# Correction of missing-wedge artifacts in filamentous tomograms by template-based constrained deconvolution

**DOI:** 10.1101/827006

**Authors:** Julio Kovacs, Jun Ha Song, Manfred Auer, Jing He, Willy Wriggers

**Affiliations:** Department of Mechanical and Aerospace Engineering, Old Dominion University, Norfolk, VA; Cell and Tissue Imaging, Molecular Biophysics and Integrated Bioimaging Division, Lawrence Berkeley National Lab, Berkeley, CA; Department of Computer Science, Old Dominion University, Norfolk, VA

## Abstract

Cryo-electron tomography maps often exhibit considerable noise and anisotropic resolution, due to the low-dose requirements and the missing wedge in Fourier space. These spurious features are visually unappealing and, more importantly, prevent an automated segmentation of geometric shapes, requiring a highly subjective, labor-intensive manual tracing. We developed a novel computational strategy for objectively denoising and correcting missing-wedge artifacts in the special but important case of repetitive basic shapes, such as filamentous structures. In this approach, we use the template and a non-negative “location map” to constrain the deconvolution scheme, allowing us to recover, to a considerable degree, the information lost in the missing wedge. We applied our method to data of actin-filament bundles of inner-ear stereocilia, which are critical in hearing transduction processes, and found a good overlap with the experimental map and with manual tracing. In addition, we demonstrate that our method can also be used for membrane detection.

## Introduction

Cryo-electron tomography (cryo-ET) of unstained frozen-hydrated samples is a widely used technique for imaging of supramolecular complexes in organelles, cells and tissues in their near-native state[1]. It consists in tilting the sample in the electron microscope to obtain a series of views of the specimen in known orientations. A disadvantage of cryo-ET when used for biological specimens is that radiation damage limits the number of images that can be obtained with this approach. This limitation in the electron dose results in typically high noise levels. Furthermore, the tilting range of the electron microscope is almost always limited, resulting in missing views (“missing wedge” in the case of single-axis tilting) and, consequently, anisotropic resolution, with that in the direction of the electron beam (*z* coordinate) showing a reduction of 50% to 66% relative to that on the *x-y* plane.

Previous approaches have addressed the problem of detecting and tracing filaments in electron tomograms by maximizing the local cross-correlation with a cylindrical template as a function of the template’s direction at each voxel[2, 3, 4]. The first two of these approaches[2, 3] use a simple compensation for the missing wedge, i.e. by stretching the template in the *z* direction, while Weber[4] uses a more accurate modeling, by convolving the template with the “resolution kernel” (inverse Fourier transform of the data-covered region in Fourier space), similarly to what we do here.

These methods yield good results in general, but encounter difficulties in cases of densely packed structures, i.e. when the map resolution is comparable to, or worse than, the spacing between filaments, due to signal blending. High noise levels typical in cryo-ET further exacerbates this issue, rendering pointwise approaches unable to yield reliable and unambiguous recovery of the underlying signal.

Here we propose a global approach to address the problem of missing wedge for such “hard” cases of extremely low SNR. Instead of sequentially scanning the image voxel wise and maximizing the correlation vs. template orientation at each voxel, we proceed in a *parallel* fashion, seeking a map *U* whose convolution with the template and the resolution kernel (mentioned above) reproduces the given experimental map (save for the noise). Such an approach could not work without additional constraints, due to the missing wedge of data in Fourier space. However, by considering the physical meaning of such convolution—namely placing copies of the template as prescribed by the map *U*— we see that the values of *U* must all be *non-negative*. Through the use of this prior knowledge, we can obtain *U* by solving a least-squares optimization problem under linear constraints.

## Results

### Modeling assumptions

The switch from sequential to simultaneous approach comes at the cost of needing to solve an extremely large system of equations. In its most general way, these equations include 3 unknowns for each voxel of the map: the coefficient of the template centered at that voxel plus the two polar angles specifying the direction of the template at that voxel. This typically means hundreds of millions of variables. Furthermore, the dependence on the polar angles is nonlinear.

For the present work we restrict ourselves to a simpler (but still useful) case of tomograms in which the direction of the pattern can be assumed to change relatively slowly across the map (Figure 1). In this case we divide the whole map into regions of small enough size that the direction of the pattern (e.g. filaments) can be considered to be nearly constant in each region. As we show later, this assumption still allows significant flexibility on the directions, even as high as about 25°deviation relative to the average direction within the region.

**Figure 1:**
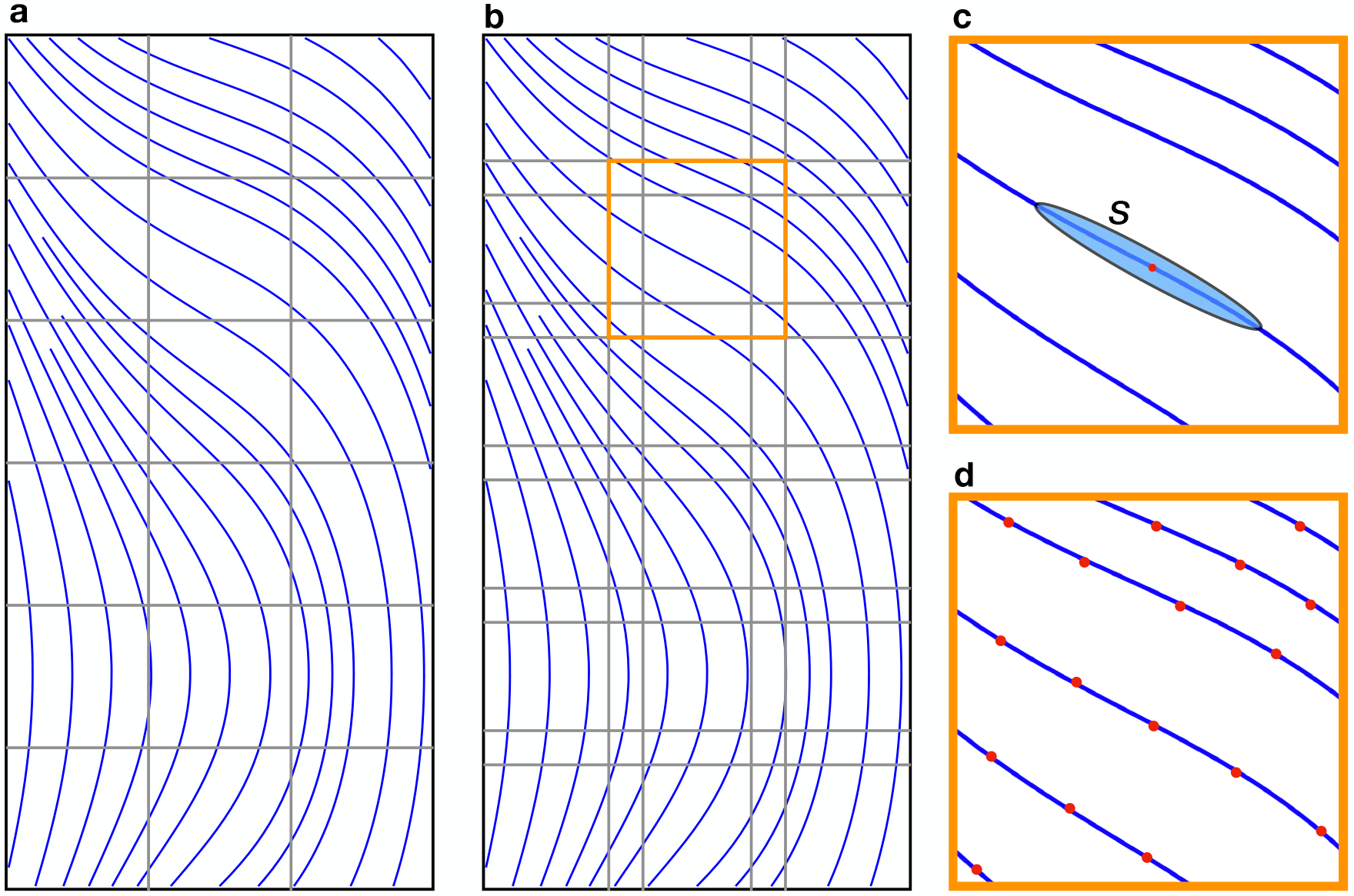
General schematic overview of our approach. **a**, the whole tomogram is partitioned in small enough regions within each of which the filaments (or the relevant repeating units) can be expected to have nearly constant direction. **b**, the partitions are enlarged to create overlap between them, in order to avoid boundary effects. **c**, focusing on one of the regions, the shape kernel *S* is shown centered at a generic location on the map, marked by a red dot. This shape kernel is represented as a Gaussian function whose longitudinal FWHM is the template length, and whose transversal FWHM is the thickness of the filament (assumed to be known). **d**, the solution of the constrained-deconvolution equations yields a map *U* that indicates the locations (depicted by red dots) where copies of *S* are to be placed (along with the corresponding scalings) to recover the “true map” (free of noise and missing-wedge artifacts), according to eq. (1).

Focusing, then, on one of these regions, the template or “shape kernel” *S* (Figure 1**c**) is assumed to have the same direction at every point, and so the convolution mentioned above is of the standard type. Thus, our main assumption is that, within each of the regions, the “true map” *T* can be expressed as:

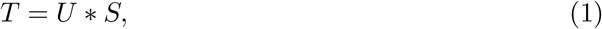

that is:

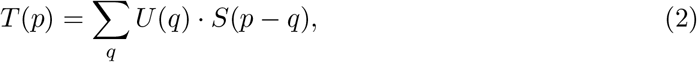

where the summation runs over all points *q* of the map.

Usually, for filament detection and tracing, the template is taken as a simple cylindrical mask of certain length and diameter. In our approach, however, it is more appropriate to use a smooth function which, for mathematical convenience, is a Gaussian function, and this is why we call it here “shape kernel” rather than template. This allows a proper formulation of the model as a kernel-based expansion given by the above equation.

Our second assumption is the image-formation model:

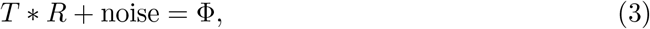

where Φ is the experimental map, and *R* is the resolution kernel, i.e. the point-spread function of the final reconstruction. The Fourier transform of *R* is the mask in Fourier space defining the data-covered region. In the case of single-axis tilt geometry, on which we are focusing in his paper, this region is a cylinder with a wedge excluded (Figure 2). Other tilt geometries are easily accommodated by using the mask corresponding to their data coverage.

**Figure 2:**
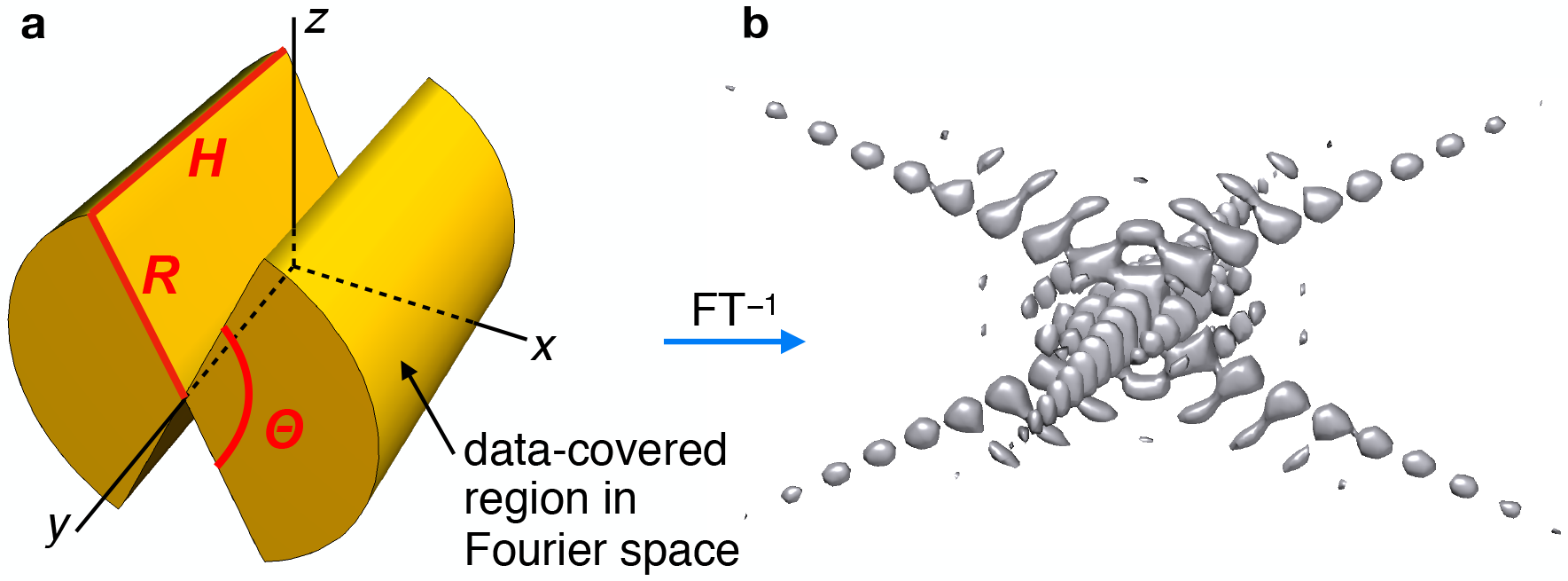
Resolution kernel. **a**, the missing wedge in Fourier space is the region where data is not available due to the limited tilt angles of the specimen (about the *y* axis). This is given by the angle Θ < 180°. Likewise, data is not available outside of the cylinder shown. These conditions limit the resolution of the reconstruction and make it worse along the *z* axis (missing-wedge direction). **b**, inverse Fourier transform of the data-covered region, depicted as an isolevel surface. This is the effective point-spread function of the tomogram, which we call here *resolution kernel*. This enters the image-formation model according to eq. (3).

### Solution of the equations by constrained deconvolution

By combining equations (1) and (3) we get:

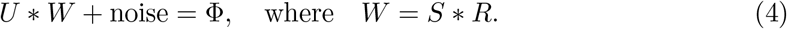

This is a *constrained deconvolution* problem, which can be recast as a least-squares problem:

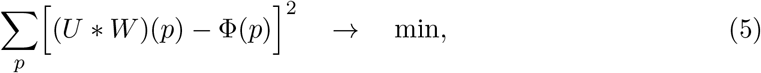

subject to the constraints *U*(*p*) ≥ 0 for all voxels *p*. This leads to the following *quadratic programming* problem:

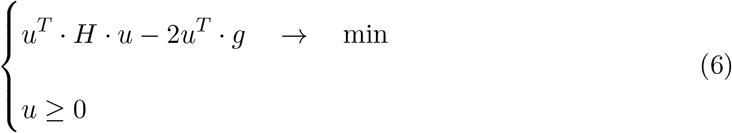

where the matrix *H* and the vector *g* involve the kernel *W*, and the vector *g* involves, in addition, the map Φ. The solution vector *u* is a 1-dimensional version of the map *U*, which will in general have relatively few voxels where it is non-zero—i.e. it is a discrete map. Once we have solved these equations for *U*, the “true map” *T* is obtained via equation 1. The maps *T* and *U* can then be used to carry out the tracing of the filaments, as described in the Methods section.

A similar constrained-deconvolution approach has been proposed years ago in the context of biopharmaceutics[5], although without the smoothing component *S*, which is essential in our case to increase the SNR. This extra factor *S* can be seen as a change of variables: instead of using *T* directly in an equation such as *T* * *R* + *N* = Φ subject to *T* ≥ 0, we use equation (4) to solve for *U* under *U* ≥ 0 and then change variables back through *T* = *U* * *S*. This provides a built-in mechanism to effectively denoise the data in an adaptive manner.

### Application to stereocilia tomograms

Stereocilia are actin-bundle-filled membrane protrusions of the apical hair cell surface. Within individual stereocilia, the precise 3D organization of the whole actin bundle has never been determined experimentally; instead, most of our knowledge is derived from transmission electron micrograph images of 70–100-nm ultrathin cross-sections directions[6]. Such studies revealed that actin filaments generally appear to be spaced ≈12 nm apart in a hexagonal pattern in the shaft region, which constitutes the bulk of the stereocilium length.

Whole-mount cryo-electron tomography data had been collected from unstained frozen-hydrated individual stereocilia[7]. The 3D volumes contain over 300 filaments per stereocilium that must be traced in their entirety. So far, manual tracing has been the only way to trace the filaments in such challenging datasets, and it is very laborious: it takes weeks for each dataset. While the manual tracing in selected intervals was key and served as a starting point for semi-automated tracing, automated approaches ensure an optimal local fit for every slice of the tomogram.

Figure 3 shows side and top views of the tomograms mentioned above. The shaft region of the actin bundle consists of mostly parallel actin filaments in a nearly hexagonal arrangement, whereas the taper region is more disorganized and hence significantly more challenging for manual as well as automated approaches. Besides the missing wedge artifacts, manifested as the complex point-spread function (Figure 2(b)), both tomograms exhibit an additional artifact, as banded patterns clearly visible on the top views. These bands are caused by the imbalance of the Fourier spots intensities, presumably due to variations in the defocus for different tilt angles (Figure 4(a)). We addressed this problem by the use of a “balancing filter” which restores the intensities of the weak spots (closer to the missing wedge), equalizing them with the strong spots (Figure 4(b)). The effect of using this type of filter is exemplified in Figure 4(c)-(e), with (c) being the top view of the same region shown in Figure 3(a), and (d)-(e) contrasting the results without (d) versus with (e) the use of the filter.

**Figure 3:**
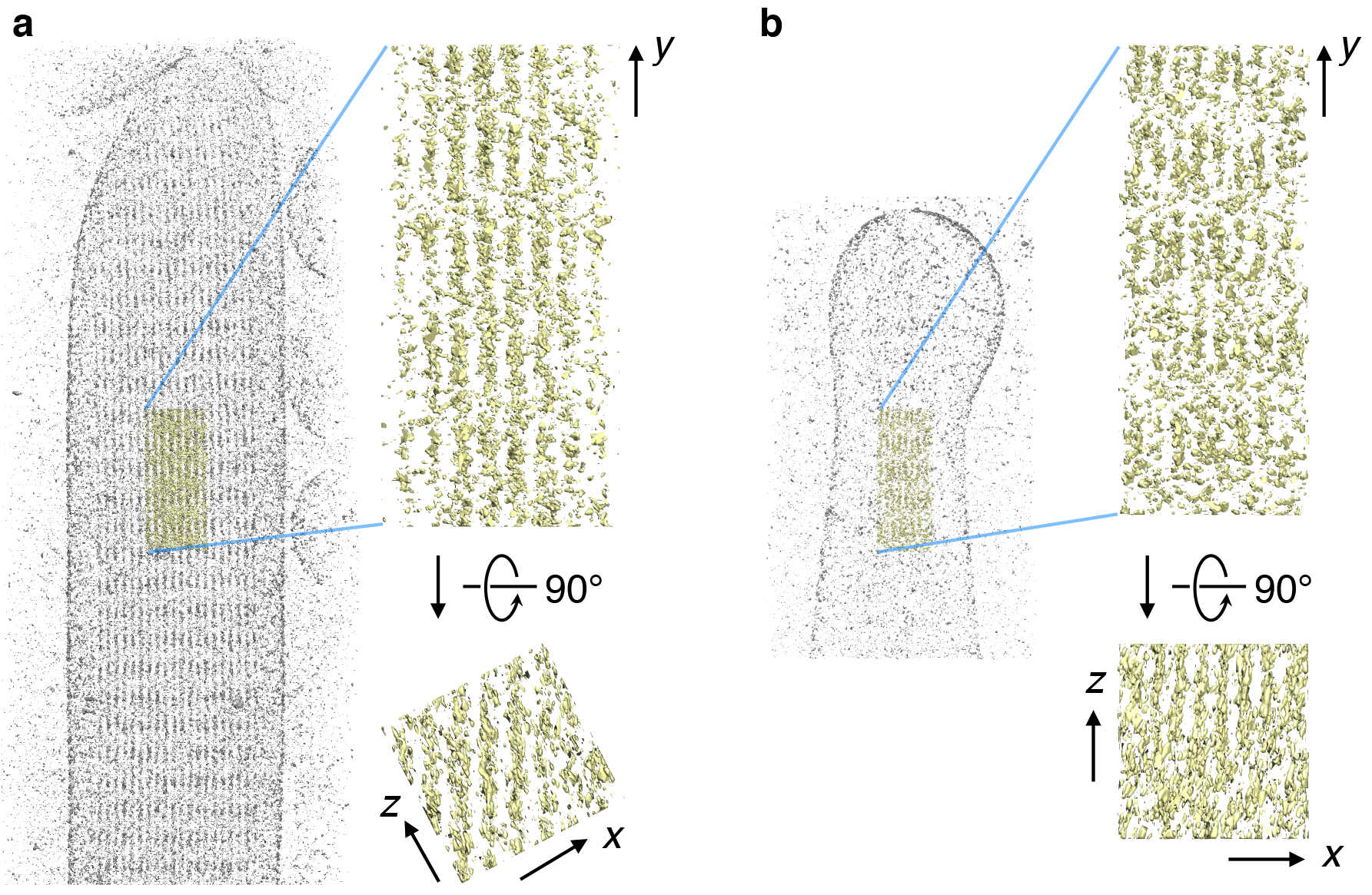
Side and top views of the two stereocilia tomograms analyzed in this paper. The yellow regions, located in the central part of each tomogram, show in more detail the artifacts that we address: noise, missing wedge, and imbalance of Fourier spot intensities. **a**, the shaft region is mostly ordered, with filaments running nearly parallel to the *y* axis in an approximate hexagonal arrangement. **b**, the taper region is less organized, with filaments exhibiting more pronounced twists and turns. The bands, clearly visible in the top views of both tomograms, are a result of the Fourier spot imbalance, presumably due to variations in defocus at different tilt angles.

**Figure 4:**
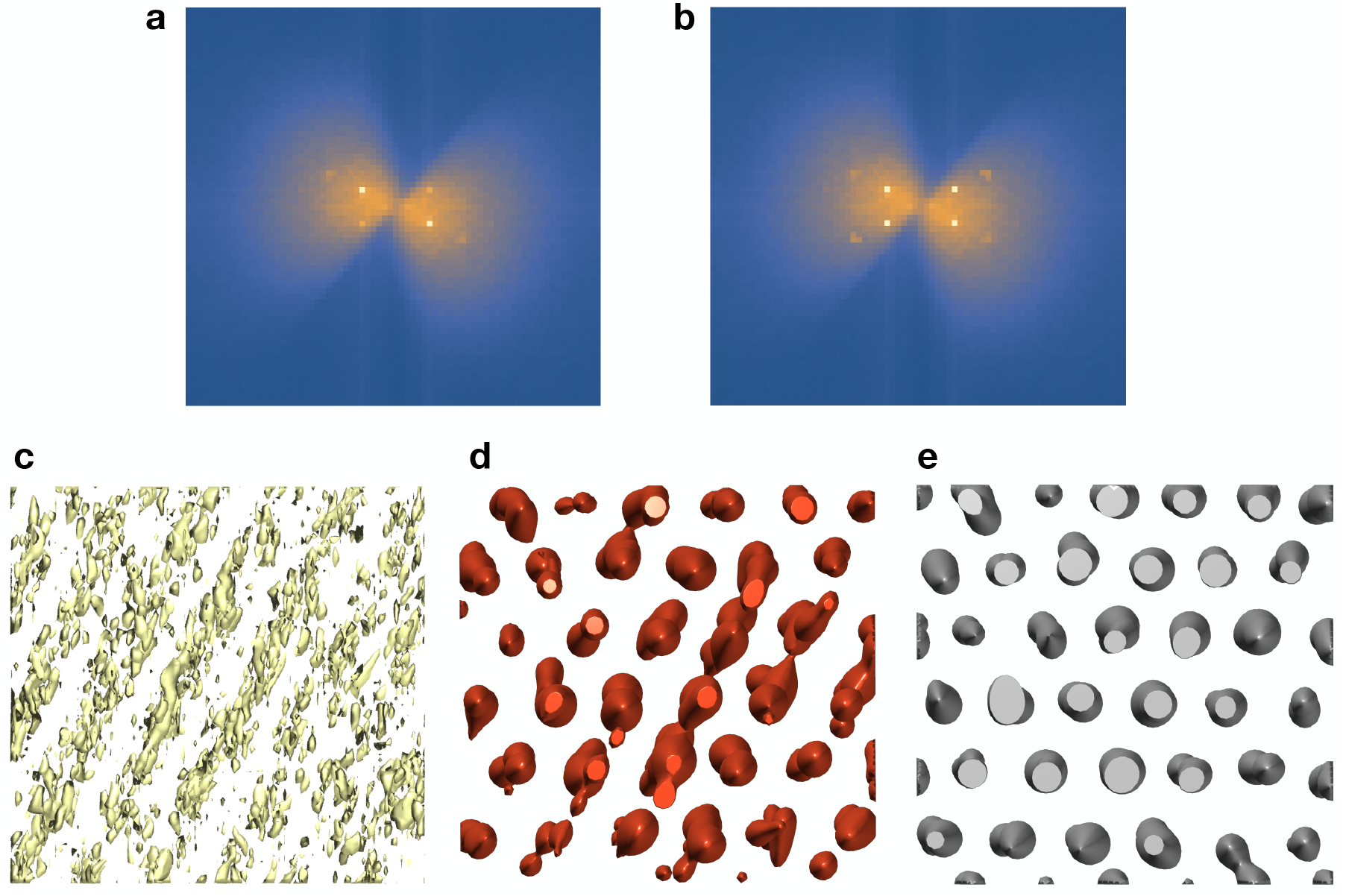
Sometimes, an imbalance among the various Fourier spots becomes apparent, as in the tomogram of the shaft region of stereocilia, with its nearly hexagonal packing of filaments. The expected power spectrum (seen down the filament direction) would have four spots of the same intensity (plus two inside the missing wedge), but the observed power spectrum shows one pair of spots with significantly weaker intensity (**a**). This results in the banded structure of the tomogram (**c**). The result of applying of approach directly shows remnants of these artifacts (**d**). By scaling the weaker spots (i.e. those nearer the missing wedge) to match the intensity of the stronger spots (**b**), we obtain a result where these artifacts are virtually eliminated (**e**).

A preliminary attempt at a semi-automatically tracing filaments in the shaft region has been proposed[8]. A set of “seed points” is manually selected by the user at one of the end faces of the bundle, and then the algorithm would trace the filaments from those starting points. The success of this approach, however, was partial, since the missing wedge artifacts made it necessary to restrict the *z* coordinate in order to obtain reasonable traces. Given the more complex organization of the taper region[9], we were concerned that this approach may not work in the taper region of the bundle.

We applied our method to both the shaft and taper regions. The results from the shaft region revealed that the filaments exhibit slight nonlinear variations in the *x* and *z* coordinates when traced along *y*, and match very well with the experimental map (Figure 5(a,b)).

**Figure 5:**
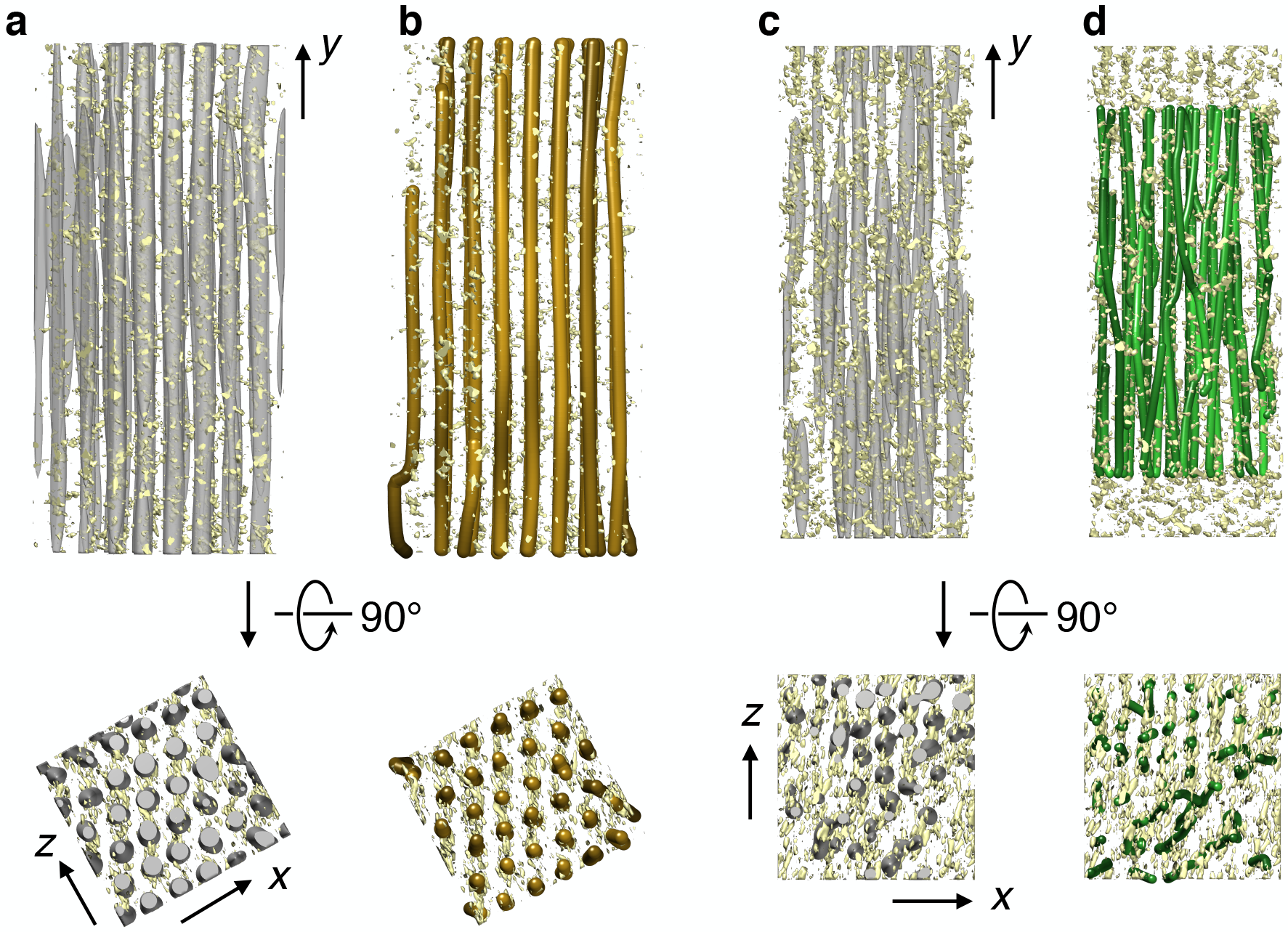
Results of our approach on the subregions shown in Figure 3, superposed on the original tomogram data. **a** and **b**, subregion of the shaft tomogram. In **a** the deconvolved map *T* is shown as a gray isocontour surface, while **b** shows the corresponding traces. Some distortions appear near the boundaries, since these results are based only on the data in these particular subregions. **c** and **d**, subregion of the taper tomogram. In **c** the deconvolved map *T* is shown as a gray isocontour surface, while **d** shows the corresponding traces. The boundary effects here are more pronounced and would prevent a correct tracing if started right at the edge in the *y* axis. Hence the tracing excludes the portions of the map within half of the length of the shape kernel. (The tracing of the global map is done in this way.) The variations in *x, z* makes it more difficult to assess the quality of the tracing from the side view, but the top view reveals a good agreement especially in the parts that are more ordered, such as the upper half of the top view.

**Figure 6:**
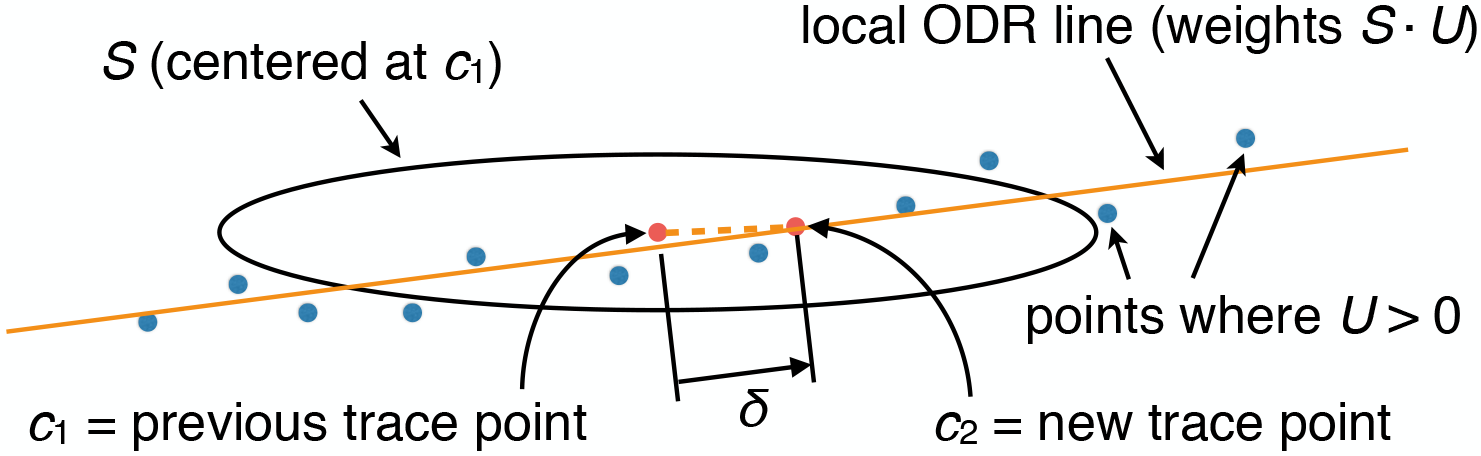
Tracing of the map *T* by means of a local regression technique. Centering the shape kernel *S* at the previous (or first) trace point *c*_1_, a local “orthogonal distance regression” line is computed based on nearby non-zero points *p* of the *U* map, using weights *S*(*p* − *c*_1_)*U*(*p*). A new trace point *c*_2_ is then taken on this line at a prescribed distance *δ* from the point on the line closest to *c*_1_.

In the taper region (Figure 5(c,d)) it is harder to compare filament traces with the experimental map due to the curvature of filaments, making the judgement of the matching accuracy more difficult than for the shaft region (where the straight and aligned filaments allow the use of projection in the common direction for a visual assessment).

Notwithstanding, a good agreement with the map density can be observed in the example cross-sections (Figure 7).

**Figure 7:**
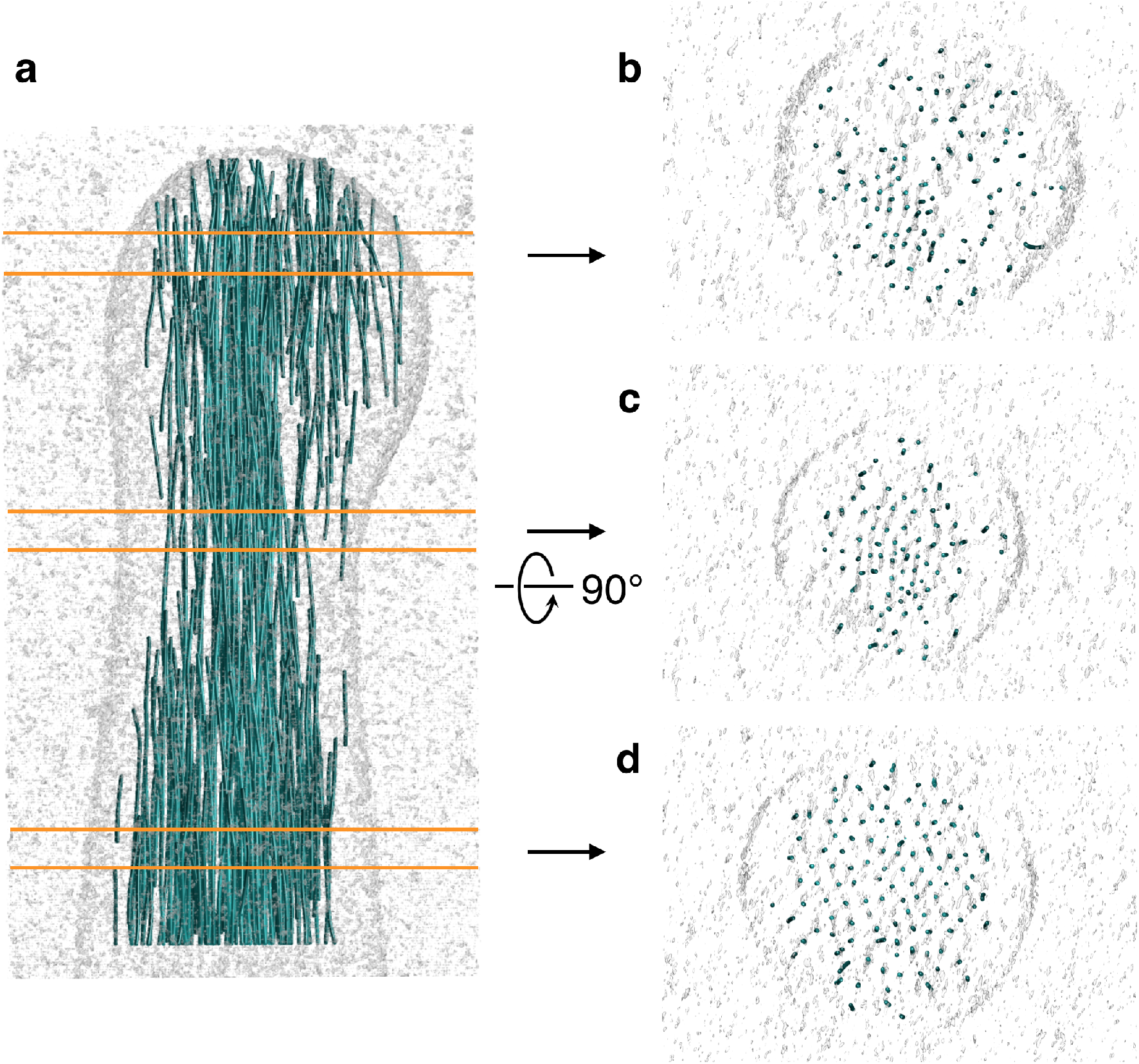
Tomogram of the taper region and top views of the three indicated slabs. Each slab is 30 voxels thick. In all of them we can discern a very good agreement between the traces and the density. The bottom slab (**d**) is the most ordered, as can be seen by its roughly hexagonal packing of the filaments. In the middle and upper slabs the filaments are less well organized, and one can see some instances of density without trace, and some of trace without density. This is because filament traces shorter than the length of the shape kernel (50 voxels in our case) were excluded, and also small gaps in density (i.e. density lower than the isocontour value used to display the map) belong to actual filamentous densities that extend above and beneath the slab.

## Discussion

We experimented with a range of lengths for the shape kernel, from 20 to 80 voxels of FWHM, and found the best compromise to be 50 voxels. This value is consistent with a previous report[2] of an optimal trade-off length of the cylindrical template of about 42 nm: 50 voxels = 50 × 0.947 nm = 47 nm).

Our approach can also detect membranes, if they have a roughly cylindrical shape in each region of the partition. Figure 8 shows traces in the membrane region detected in the stereocilia taper tomogram. The parts of the membrane (the “neck” in case) whose direction is more than about 25° from the average filament direction could not be detected, as the attenuation at those angles is higher than the prescribed detection threshold. This property of detecting membranes can be helpful and complement other methods more specifically targeted at membrane detection.

**Figure 8:**
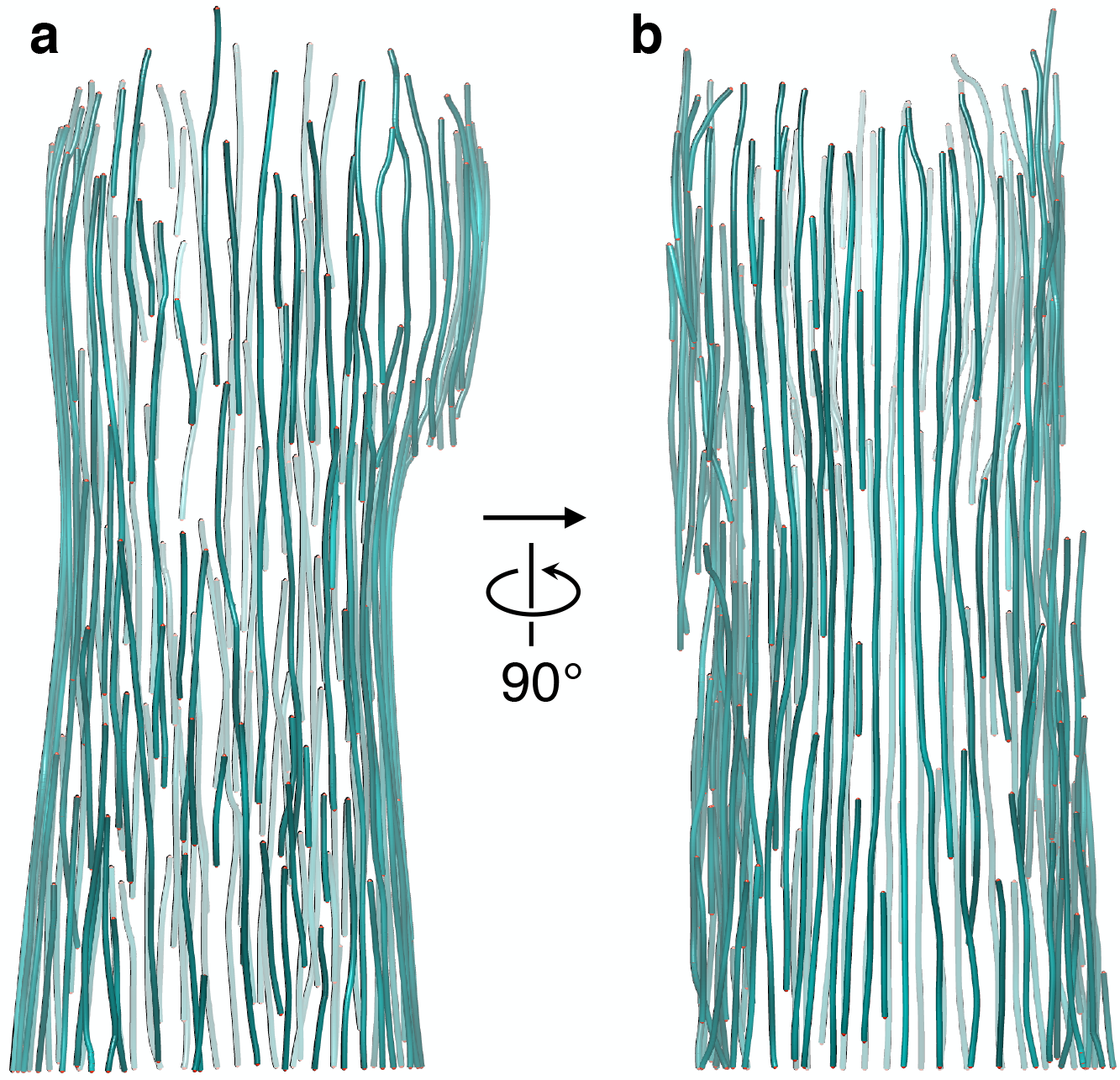
Side views of the traces in the membrane region of the taper sterocilia tomogram. Membranes are detected as they are seen by the algorithm as arrangements of 1D paths. This property can be helpful and complement other methods more specifically targeted at membrane detection.

In cases where subtomogram averaging is not helpful to reduce missing-wedge artifacts due to the parallelism of the various copies of the structure, our method becomes an excellent complementary alternative, since it works best when the repeating subunits are nearly parallel to one another.

Future developments we are planning include:

- Extension to other tilting geometries, such as double-axis tilt and conical tilt, which involve a modification in the resolution kernel *R*.
- Increased ability to handle multiple and highly variable filament directions, by adding two extra variables at each voxel (describing the template direction vector at the point). This will require the development of novel methods to solve the 3 × larger (and nonlinear) system of equations.
- As a particular case of the above, ability to detect membranes in arbitrary shape and direction.

## Methods

### Cryo-Tomography of Inner Ear Sensory Epithelial Hair Cell Stereocilia

We used cryo-tomogram data sets of stereocilia that existed in the Auer lab and have been described extensively elsewhere[7].

### Constrained deconvolution equations lead to a quadratic programming problem

Here we write down the equations in the general case where a variable direction of the shape kernel *S* is allowed. Let us denote this dependence by *S_y_*. Then, instead of the special main assumption *T* = *U* * *S* (eq. 1) we have the following general one:

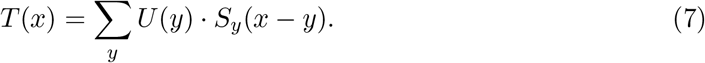

The image-formation model (eq. 3) remains unchanged. Writing the convolution explicitly:

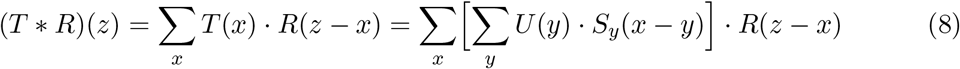

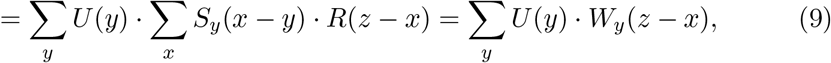

where

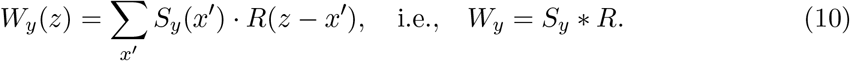

Hence the general least-squares equations for *U* are:

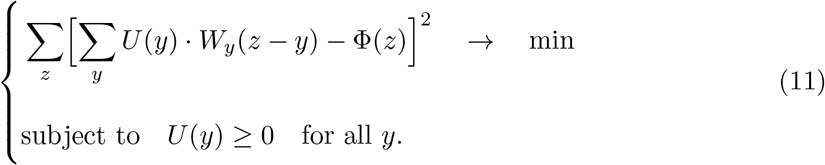

Note that if the shape kernel direction is the same for all points in the map, then the *W* kernel is too: *W* = *S* * *R*, as in eq. (4), and we obtain eq. (5) as a special case of eq. (11).

If the tomogram is of sufficiently good quality that it allows the determination of the shape kernel direction at each point beforehand, as done in previous works[2, 3, 4], then our approach would not be needed. In hard cases like the stereocilia data described in the Results section, attempts at prior determination of the filament direction at each point utterly failed due to the high levels of noise. This is the very reason that a global approach like our is imperative for this type of tomographic data.

Expanding the quadratic form:

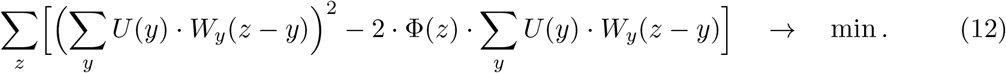

If *N* = number of voxels in the map, and the voxels are enumerated as {*x_i_*}_*i*=1,…,*N*_, we define the *deconvolution matrix A* by:

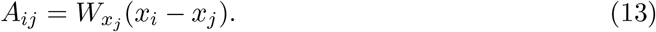

Let us also use the notation *U_i_* = *U*(*x_i_*) and Φ_*i*_ = Φ(*x_i_*). Then our least-squares problem takes the form:

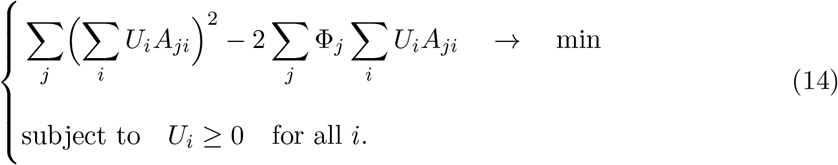

In order to write the equations in a matrix-vector notation, let us use the following column-array representation of the maps involved: *u* = (*U*_1_,…, *U_N_*)^*T*^ and *f* = (Φ_1_,…, Φ_*N*_)^*T*^, where the superscript *T* denotes the transpose. Then Σ_*i*_ *u_i_A_ji_* = (*Au*)_*j*_ and so:

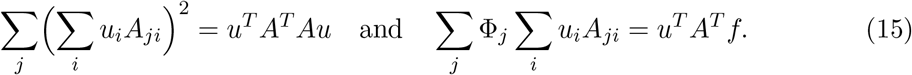

Therefore, denoting *H* = *A^T^ A* and *g* = *A^T^ f*, we arrive at our quadratic programming problem:

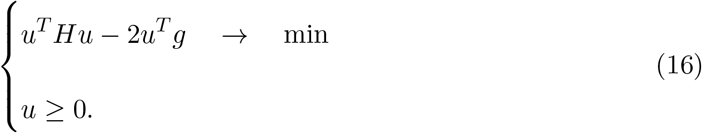

As pointed out earlier, these equations involve, in general, 3*N* unknowns: *N* coefficients *u_i_* and 2*N* parameters describing the direction vector of the shape kernel at each of the *N* voxels. The dependence of these equations on *u* is quadratic, but the dependence on the direction vectors (on which *H* and *g* depend) is more complex. We envision the development of more sophisticated methods to solve such systems of equations under these general conditions. As mentioned before, for the purpose of the present paper, we assume that the whole tomogram can be partitioned in regions within each of which the filament direction can be considered approximately constant. In this case the matrix *H* and the vector *g* are constant within each region. Each subsystem then involves only *N_k_* + 2 unknowns instead of 3*N*, where *N_k_* is the number of voxels in region *k*. The various regions should overlap along their boundaries, in order to avoid aliasing and boundary effects (Figure 1).

### Solution of the quadratic programming problem

We solved eq. (16) iteratively within each of the regions. We start by using the initial direction of the shape kernel as the *y* axis. With this direction fixed, we solved eq. (16) by means of the software package SNOPT[10]. This step was wrapped inside a loop consisting of a Newton-Raphson scheme to minimize the objective function with respect to the direction vector of the shape kernel (that is, the remaining 2 variables, on which *H* and *g* depend). This process converged very quickly, in two or three iterations. In this manner we obtained the *U* maps for all the regions. Each of these maps was then cropped along its boundaries by half of the overlap width, yielding a global *U* map corresponding to the whole tomogram. The global *T* map was then computed by means of eq. (7).

### Filament tracing using the global *U* and *T* maps

In principle, tracing could be done by just using *T*, but the availability of *U* allows for a much simpler and faster procedure.

After obtaining the global *U* and *T* maps as described in the previous section, the voxels of the *T* map are scanned sequentially in order of increasing *y* coordinates (Figure 6). Each voxel is checked to determine if it is a local maximum of *T* on that particular *x-z* cross-section. If so, a new trace starts from this voxel *c*_1_. To obtain the next trace point, a local “orthogonal distance regression” is performed based on the *U* map: the shape kernel *S* is centered at *c*_1_, and the voxels *p* where *U*(*p*) > 0 are weighted according to *S*(*p* − *c*_1_) · *U*(*p*). These weights are used in the regression, which yields a line that best fits those points *p* in a local neighborhood of *c*_1_. Denoting by *c*′ the point on the regression line closest to *c*_1_, the next point *c*_2_ of the trace is taken at a prescribed distance *δ* from *c*′. When a trace point is reached whose *T* value is less than a prescribed threshold, the trace ends. Every trace point determines a ball of “used” voxels around it, which serve to avoid repetitions of starting voxels and collisions with previously determined traces.

### Balancing of Fourier spot intensities

In the tomogram of the shaft region of the tomogram, which exhibits a clear hexagonal pattern in the bundle, we noticed an imbalance of the Fourier peaks relative to the expected spectrum of a hexagonally symmetric structure Figure 4. This seems to be due to a variation in the defocus of the microscope when the tilt angle changes. Hence, we used the hexagonally symmetric shaft portion to define an ad-hoc “balancing filter”, which equalizes the magnitudes of the weak spots (located nearer the missing wedge) to those of the corresponding strong spots (located at 60°from the weak ones). This filter *F* was introduced in the image-formation model thus:

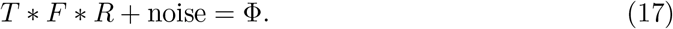

Our method then proceeds as described above, but with *R* replaced by *R*′ = *F* * *R*.

## References

[1] Irobalieva, R. N., Martins, B. & Medalia, O. Cellular structural biology as revealed by cryo-electron tomography. Journal of Cell Science 129, 469–476 (2016). URL http://jcs.biologists.org/content/129/3/469.

[2] Rigort, A. et al. Automated segmentation of electron tomograms for a quantitative description of actin filament networks. Journal of Structural Biology 177, 135–144 (2012). URL http://www.sciencedirect.com/science/article/pii/S1047847711002516.

[3] Rusu, M., Starosolski, Z., Wahle, M., Rigort, A. & Wriggers, W. Automated tracing of filaments in 3D electron tomography reconstructions using Sculptor and Situs. Journal of Structural Biology 178, 121–128 (2012). URL http://www.sciencedirect.com/science/article/pii/S104784771200069X. Special Issue: Electron Tomography.

[4] Weber, B. et al. Automated tracing of microtubules in electron tomograms of plastic embedded samples of Caenorhabditis elegans embryos. Journal of Structural Biology 178, 129–138 (2012). URL http://www.sciencedirect.com/science/article/pii/S1047847711003509. Special Issue: Electron Tomography.

[5] Verotta, D. An inequality-constrained least-squares deconvolution method. Journal of Pharmacokinetics and Biopharmaceutics 17, 269–289 (1989). URL https://doi.org/10.1007/BF01059031.

[6] Shin, J.-B. et al. Molecular architecture of the chick vestibular hair bundle. Nature Neuroscience 16, 365–374 (2013). URL http://dx.doi.org/10.1038/nn.3312.

[7] Metlagel, Z. et al. Electron cryo-tomography of vestibular hair-cell stereocilia. Journal of Structural Biology 206, 149–155 (2019).

[8] Sazzed, S. et al. Tracing actin filament bundles in three-dimensional electron tomography density maps of hair cell stereocilia. Molecules 23 (2018). URL http://www.mdpi.com/1420-3049/23/4/882.

[9] Song, J. et al. A cryo-tomography-based volumetric model of the actin core of mouse vestibular hair cell stereocilia lacking plastin 1. bioRxiv (2019). URL https://www.biorxiv.org/content/early/2019/11/01/825737. https://www.biorxiv.org/content/early/2019/11/01/825737.full.pdf.

[10] Gill, P. E., Murray, W. & Saunders, M. A. SNOPT: An SQP algorithm for large-scale constrained optimization. SIAM Rev. 47, 99–131 (2005).

